# Complex feline disease mapping using a dense genotyping array

**DOI:** 10.1101/2021.08.09.455727

**Authors:** Isabel Hernandez, Jessica J. Hayward, Jeff A. Brockman, Michelle E. White, Lara Mouttham, Elizabeth A. Wilcox, Susan Garrison, Marta G. Castelhano, John P. Loftus, Filipe Espinheira Gomes, Cheryl Balkman, Marjory B. Brooks, Nadine Fiani, Marnin Forman, Tom Kern, Bruce Kornreich, Eric Ledbetter, Santiago Peralta, Angela M. Struble, Lisa Caligiuri, Elizabeth Corey, Lin Lin, Julie Jordan, Danny Sack, Adam R. Boyko, Leslie A. Lyons, Rory J. Todhunter

**Affiliations:** Department of Clinical Sciences, College of Veterinary Medicine, Cornell University, Ithaca, New York, United States of America; Department of Biomedical Sciences, College of Veterinary Medicine, Cornell University, Ithaca, New York, United States of America; Pet Nutrition Center, Hill’s Pet Nutrition, Topeka, Kansas, United States of America; Bioinformatics and Integrative Biology, University of Massachusetts Medical School, Worcester, Massachusetts, United States of America and Vertebrate Genomics Broad Institute of MIT and Harvard, Cambridge, Massachusetts, United States of America; Cornell Veterinary Biobank, College of Veterinary Medicine, Cornell University, Ithaca, New York, United States of America; Department of Population Medicine and Diagnostic Services, College of Veterinary Medicine, Cornell University, Ithaca, New York, United States of America; Cornell University Veterinary Specialists, Stamford, Connecticut, United States of America; Department of Veterinary Medicine & Surgery, College of Veterinary Medicine, University of Missouri, Columbia, Missouri, United States of America

**Author notes:** These authors contributed equally to this work (Isabel Hernandez, Jessica J. Hayward).

**Keywords:** feline complex disease, genome-wide association study, biobank

## Abstract

The current feline genotyping array of 63k single nucleotide polymorphisms has proven its utility within breeds, and its use has led to the identification of variants associated with Mendelian traits in purebred cats. However, compared to single gene disorders, association studies of complex diseases, especially with the inclusion of random bred cats with relatively low linkage disequilibrium, require a denser genotyping array and an increased sample size to provide statistically significant associations. Here, we undertook a multi-breed study of 1,122 cats, most of which were admitted and phenotyped for nine common complex feline diseases at the Cornell University Hospital for Animals. Using a proprietary 340k single nucleotide polymorphism mapping array, we identified significant genome-wide associations with hyperthyroidism, diabetes mellitus, and eosinophilic keratoconjunctivitis. These results provide genomic locations for variant discovery and candidate gene screening for these important complex feline diseases, which are relevant not only to feline health, but also to the development of disease models for comparative studies.

## Introduction

There are 365 hereditary disorders of cats listed on OMIA (Online Mendelian Inheritance in Animals, https://omia.org/home/ accessed June 7th, 2021), of which only 119 (32.6%) are Mendelian traits and only 136 (37.3%) have likely causal variants. Clearly, there are a large number of feline diseases whose genetic basis is still unknown. Moreover, 230 of these hereditary feline disorders are potentially good models for human disease.

Random bred cats are the most common cats in American households, accounting for 84% of the population in the United States [1]. Random bred cats comprised 89% of cats admitted the Cornell University Hospital for Animals (CUHA) in the last 15 years, thus providing an important spontaneous source of DNA for increasing sample sizes of genetic mapping studies.

Compared to purebreds, random bred cats have shorter linkage disequilibrium, due to the large number of generations since the origin of the random bred cat population, with archaeological evidence of a human and cat burial site as old as 9,500 years [2]. The genetic heterogeneity of random bred cats, the additive effect of many genes, and their environmental interaction makes discovering variants contributing to complex diseases more challenging than for Mendelian traits [3]. At least a few Mendelian traits have been mapped in random bred cats, including spongy encephalopathy, Glanzmann thrombasthenia, and inflammatory linear verrucous epidermal nevus [4–6]. Additional factors that make the discovery of complex disease genetic mechanisms difficult include sample size, phenotyping accuracy, mapping array marker density, and access to whole genome sequences for variant discovery [7].

The current 63k Illumina feline single nucleotide polymorphism (SNP) mapping array has been used successfully within breeds to map variants for Mendelian diseases. Examples include the discovery of the *WNK4* variant that causes hypokalemia in Burmese cats [8], a region on chromosome E1 associated with progressive retinal atrophy in Persian cats [9], a causal variant in *COLQ* for hereditary myopathy in Devon Rex and Sphynx cats [10], refinement of the region on chromosome B4 associated with craniofacial structure and frontonasal dysplasia in Burmese cats [11], a region on chromosome A3 associated with an inherited neurologic syndrome in a family of Oriental cats [12], and a dominant channelopathy variant causing osteochondrodysplasia in Scottish Fold cats [13]. This array has also been used in a limited number of within-breed genome wide association studies (GWAS) for complex disease [14,15], but there are no reports of GWAS performed with an across-breed design.

Here, we genotyped 1,122 cats using a one-time proprietary Illumina high density 340k SNP mapping array designed by Hill’s Pet Nutrition, in an effort to identify genetic underpinnings for nine complex diseases. Our samples consisted of a mix of 31 purebreds and 905 random bred cats, the majority of which were domestic shorthairs. This array improves upon the density of the current commercial 63k array by a factor of >5. As quality control and to validate the accuracy of the 340k array, we performed a GWAS for the *Orange* coat color locus and for Factor XII deficiency, which are known to be associated with a region on chromosomes X and A1, respectively [16–19].

The complex diseases included in this study were hypertrophic cardiomyopathy (HCM), hyperthyroidism, diabetes mellitus (DM), chronic kidney disease (CKD), chronic enteropathy, inflammatory bowel disease (IBD), small cell alimentary lymphoma (SCAL), hypercalcemia, and feline eosinophilic keratoconjunctivitis (FEK). These diseases are among the most common complex diseases of cats admitted to CUHA and are some of the most common and important feline diseases in clinical veterinary practice [20].

We used both a linear mixed model (LMM) and a multi-locus method called Fixed and random model Circulating Probability Unification (FarmCPU) to perform GWAS, and together identified loci significantly associated with hyperthyroidism, DM, FEK, and IBD. Additionally, we identified suggestive loci for the diseases HCM and hypercalcemia. Here, we describe the largest genetic mapping study of feline complex diseases with the densest mapping array ever performed.

## Results

### Validation of array

Principal component analysis (PCA) was performed using all genotyped cats that passed quality control, and showed that there was no batch effect due to the 11 sequential plates used for genotyping (Fig. 1A). The first two components, principal component (PC)1 and PC2, explained 31.3% of the total genetic variation. The cluster that separates on PC2 in this PCA includes 40 cats from a closed colony of domestic shorthair (DSH) cats from a local breeding facility, genotyped mainly on plates 7 and 11. Principal component analysis of the genotypes of all 221 purebred cats showed that PC1 separates western breeds, like Manx and Persian, from eastern breeds, like Tonkinese and Burmese (Fig. 1B). This eastern-western distribution of breeds is also seen on PC1 of the PCA of all cats (Fig. 1A) and has been shown previously using the 63k genotyping array [16,21–23]. PC2 of the purebred cat PCA separates the Devon Rex cats from the other breeds. The first two components of the purebred cat PCA explained only 16.4% of the total genetic variation, much less than the 38.4% explained by the first two components of a PCA using the 63k array [23].

**Figure 1.**
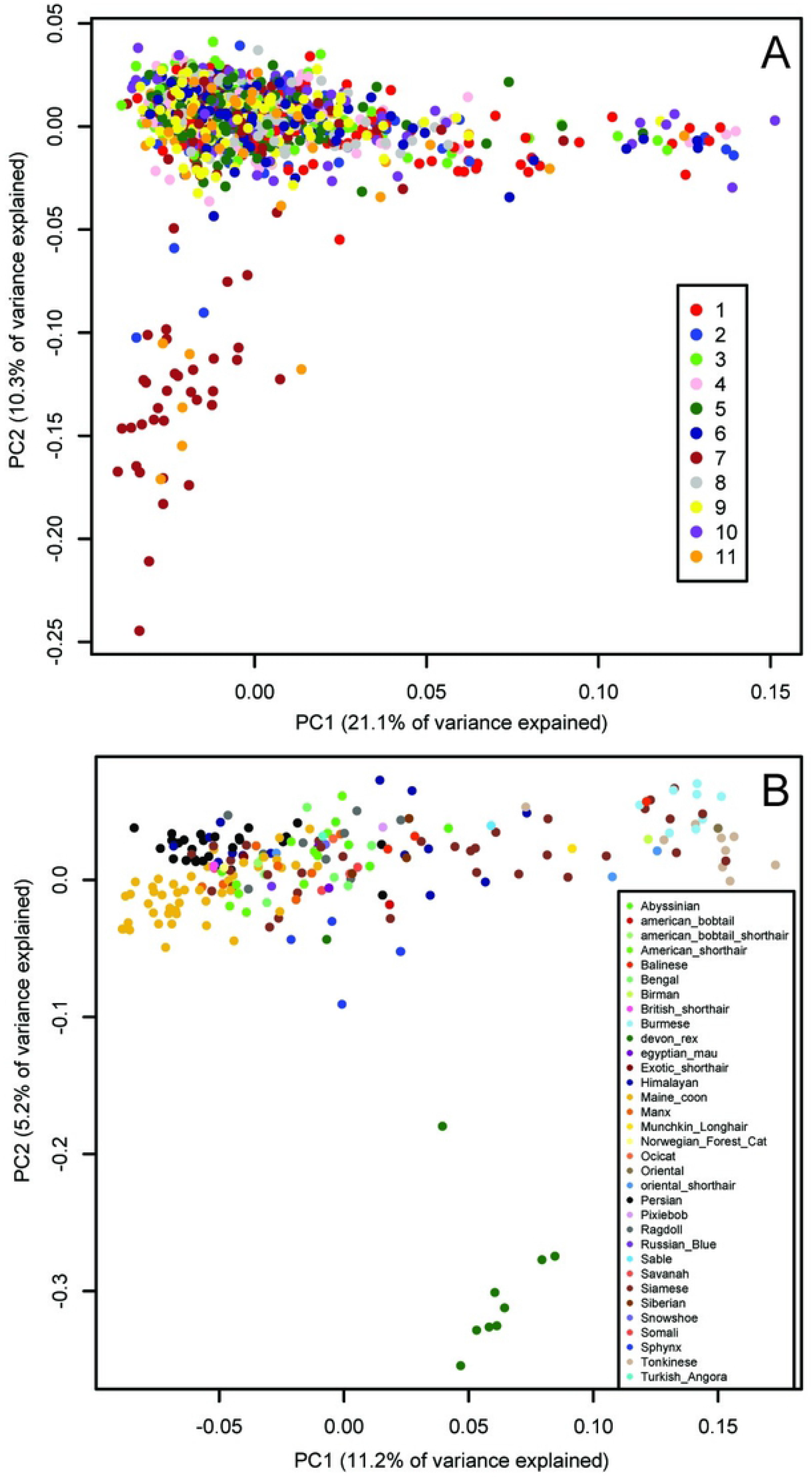
Principal Component Analysis of cat genetic structure. Dimensions PC1 and PC2 are shown. (A) All 1,122 cats that passed QC, color-coded by genotyping plate (1 to 11), showing the absence of a batch effect. PC1 shows the eastern-western breed distribution. The cluster of cats that separate on PC2 is from a local colony that were genotyped on plates 2 (dark blue), 7 (brown), and 11 (orange). (B) 221 purebred cats color-coded by breed, showing the eastern-western breed distribution on PC1. The Devon Rex breed (dark green) separates on PC2.

### GWAS positive controls

As a positive control, we performed a GWAS on the presence of orange fur in random bred cats (90 orange fur, 121 black/brown fur). Using the linear mixed model in GEMMA, we identified 25 significant associations on a region of chromosome X between 102,884,842 bp and 112,136,902 bp (Fig. 2A; S1 Table). The most significantly associated SNP in both the LMM and FarmCPU GWAS is at 110,230,748 bp (*P*=1.8×10^−102^ and *P*=2.2×10^−97^, respectively), located within an intron in the gene *Ecto-NOX Disulfide-Thiol Exchanger 2 (ENOX2).* This region is known to contain the *Orange* cat coloration locus [16–18] and the most significant SNP is within the 1.5 Mb haplotype block identified by Gandolfi *et al.* (2018). A linkage disequilibrium (LD) plot of this region showed that the 340k array has very few markers between 105-110 Mb on chromosome X, and only 4 markers within the 1.5 Mb haplotype block remain after minor allele frequency (MAF) and missingness filters, preventing the refinement of this region (S1A Fig.).

**Figure 2.**
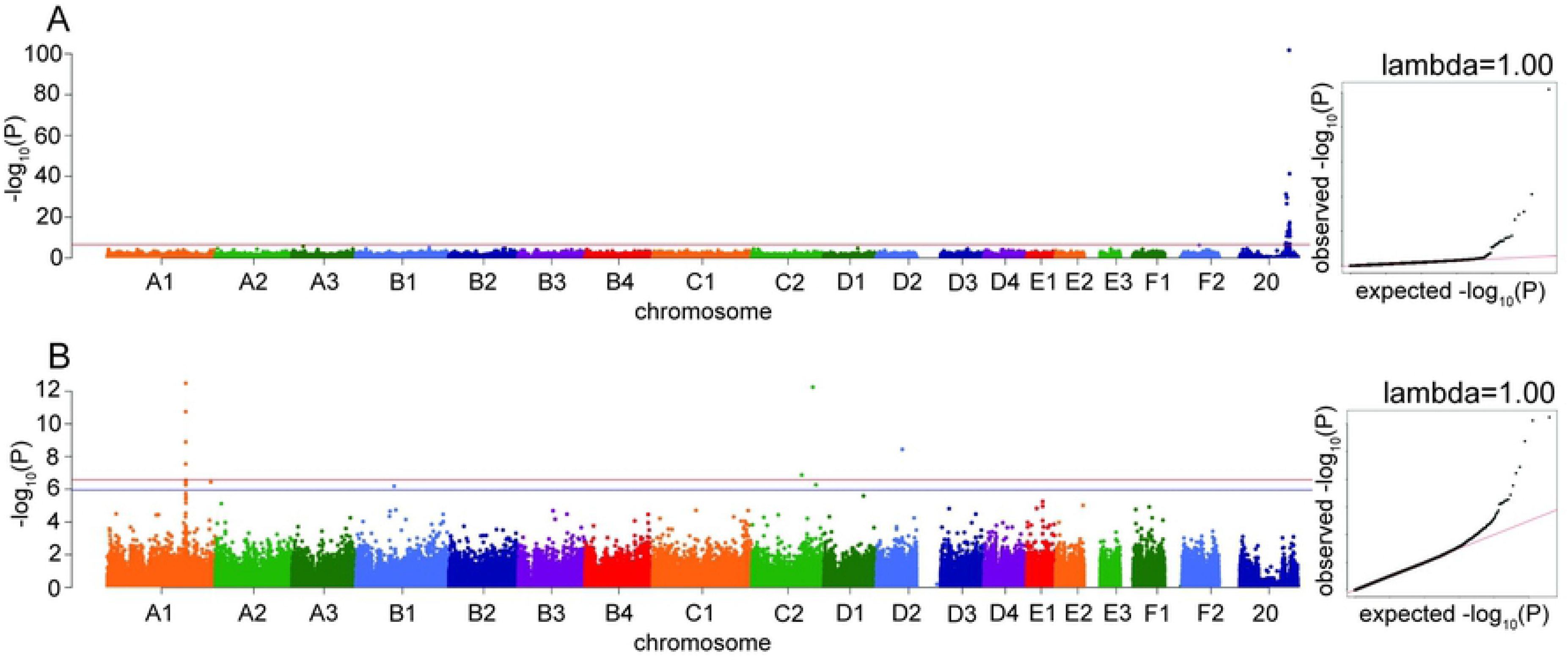
Manhattan and quantile-quantile (QQ) plots for GWAS positive controls. X axis represents the chromosomal SNP position and Y axis represents the −log_10_(*P*-value). The QQ plots show observed versus expected *P*-values for each SNP. (A) *Orange* coat color locus, showing the significant association on chromosome X (*P*=1.8×10^−102^). (B) Factor XII deficiency, showing the significant associations on chromosomes A1 and C2. The red line on the Manhattan plots shows the Bonferroni-corrected significance threshold and the blue line on the Manhattan plot in B shows the Bonferroni-corrected significance threshold calculated using unlinked SNPs. The genomic inflation factor (***λ***) is shown on each QQ plot.

As a second positive disease control, we performed a GWAS for factor XII deficiency, using 19 affecteds and 34 controls. The LMM in GEMMA identified four significant associations on chromosome A1, between 175,333,103 bp and 175,445,463 bp, which reside within 63 kb of the gene *Coagulation Factor XII (F12)* (Fig. 2B, S1B Fig.). The most significant association using the FarmCPU method was the same A1 association at 175,445,463 bp (*P*=1.4×10-19). Two high-frequency variants in the gene *F12* have previously been reported in cats with factor XII deficiency [19]. Other significant associations were also identified in the factor XII GWAS by both models, on chromosomes C2, C1, D2, F1 and D3.

### Disease GWAS

Across-breed case/control GWAS was conducted for the diseases HCM, hyperthyroidism, DM, CKD, chronic enteropathy, IBD, SCAL, FEK, hypercalcemia, and all gastrointestinal phenotypes (chronic enteropathy, IBD, and SCAL) merged together. Significance thresholds were calculated using the Bonferroni correction on all SNPs included in each GWAS, while suggestive thresholds were calculated using the Bonferroni correction on a pruned set of unlinked SNPs.

Three significant and two suggestive associations were identified above the genome-wide thresholds by the LMM GWAS in GEMMA (Table 1). The FarmCPU GWAS showed very similar results to the LMM GWAS, with significant associations for DM and hyperthyroidism (Table 1; S2 Table). However, the FEK association was not significant in the FarmCPU GWAS while the IBD association was significant (Table 1; S2 Table). Since the results from the two methods were so similar, we have chosen to focus illustrating the results of the LMM GWAS. Genomic inflation factors, ◻ are all <1.07 (range of 0.997-1.052, average 1.016 for LMM; range of 1.013-1.062, average 1.033 for FarmCPU), showing successful control for underlying population structure.

**Table 1:**
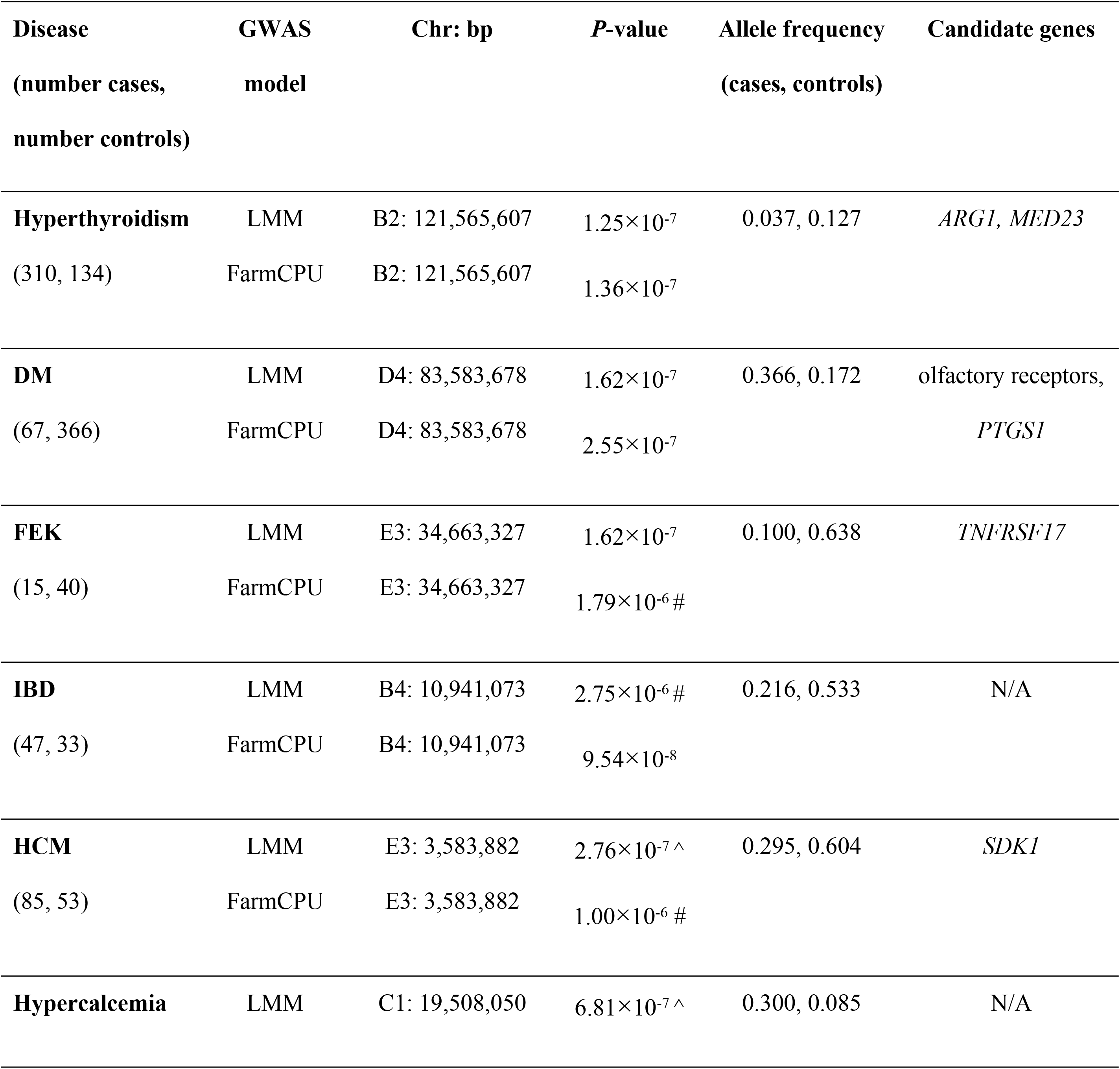
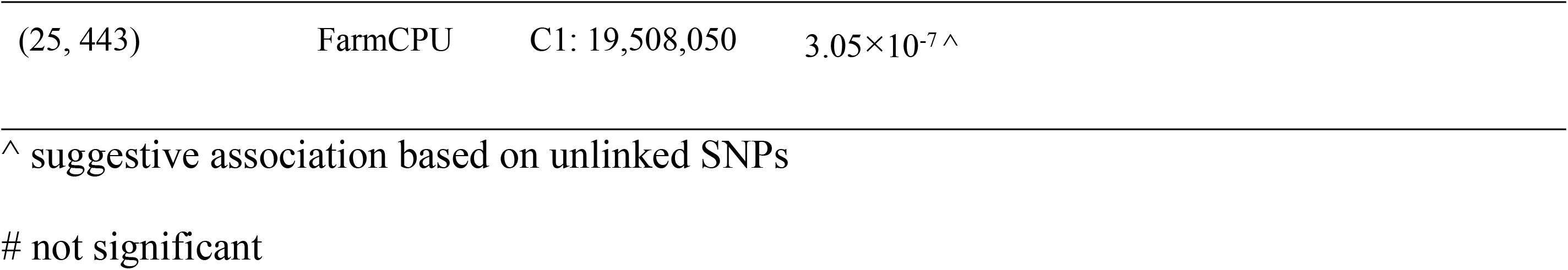
Significant and suggestive associations identified for complex diseases using an across-breed GWAS design. Results are shown for both the LMM and FarmCPU GWAS.

#### Hyperthyroidism

For hyperthyroidism, we found a solitary significant association on chromosome B2 (*P*=1.25×10^−7^ in LMM, *P*=1.36×10^−7^ in FarmCPU), located in the gene *Arginase 1 (ARG1)* and 5.5 kb downstream of, although not in LD with, the gene *Mediator Complex Subunit 23 (MED23)* (Fig. 3A). The B2 locus increases hyperthyroidism risk in DSH cats (S3 Table).

**Figure 3.**
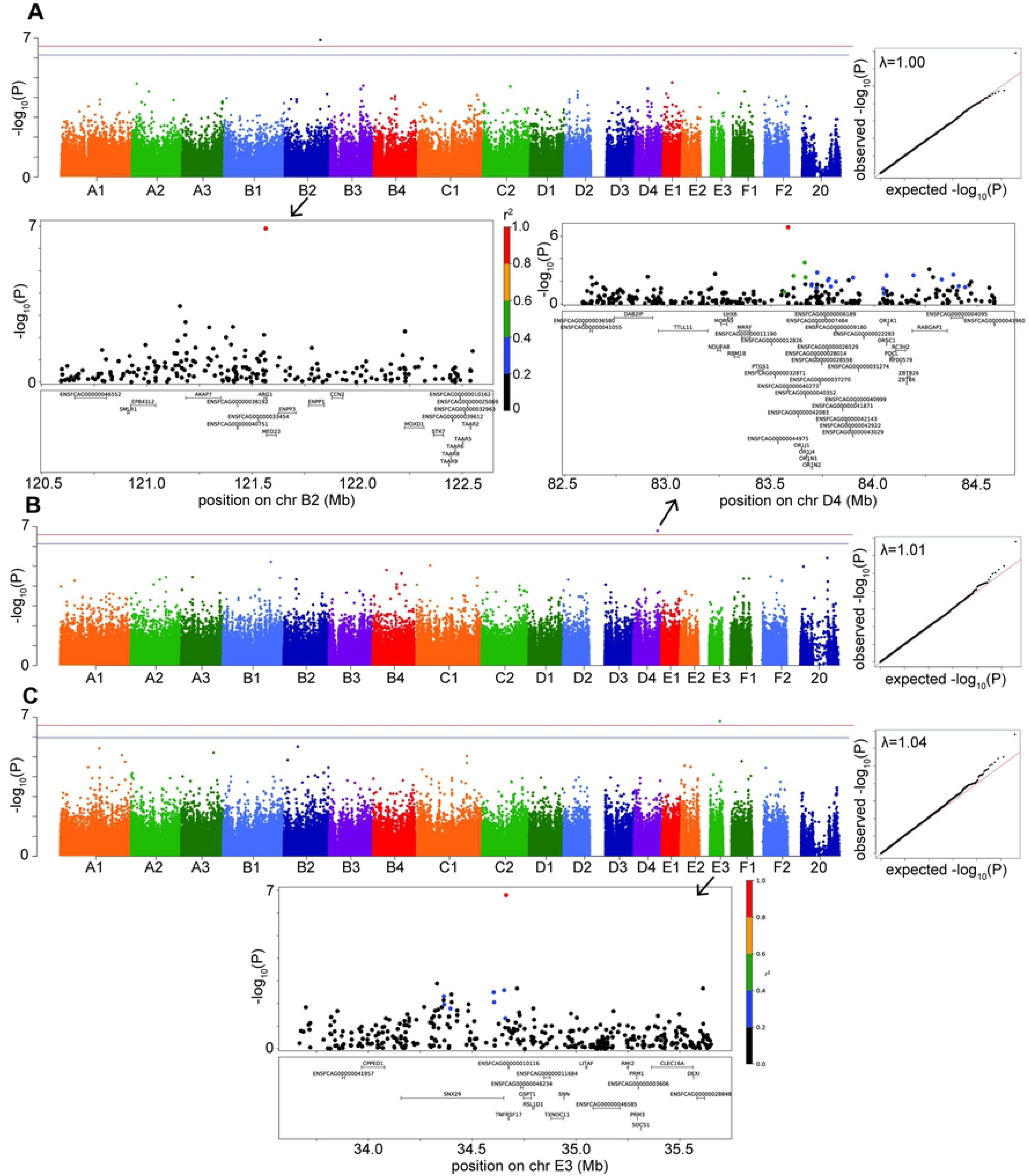
Manhattan, quantile-quantile (QQ), and LD plots for case-control disease significant associations, using the LMM GWAS results. X axis represents the chromosomal SNP position and Y axis represents the −log_10_(*P*-value). The QQ plots show observed versus expected *P*-values for each SNP. (A) Hyperthyroidism, showing the significant association on chr B2. (B) DM, showing the significant association on chr D4. (C) FEK, showing the significant association on chr E3. On Manhattan plots, the red line is Bonferroni-corrected significance threshold, and the blue line is Bonferroni-corrected significance threshold calculated using unlinked SNPs. Inflation factors (***λ***) are shown on QQ plots. On LD plots, the colors indicate the amount of LD (r^2^) with the most significant SNP, ranging from black (r^2^<0.2) to red (r^2^>0.8).

#### Diabetes mellitus

Diabetes mellitus was significantly associated with a SNP on chromosome D4 (*P=*1.62×10^−7^ in LMM, *P*=2.55×10^−7^ in FarmCPU) (Fig. 3B). The LD region includes many members of the olfactory receptor gene family, such as *OR1J*, *OR1N*, *OR1K*, and *OR5C*, among other genes. The gene *PTGS1*, (*prostaglandin synthase G/H isoform 1),* also known as *COX1* (*cyclooxygenase-1*), is located within 123 kb downstream of, although not in LD with, our significant association. This locus on D4 affects the risk of DM in DSH and Maine Coon cats, but not in cats of other breeds and DLH cats (S3 Table).

#### Feline eosinophilic keratoconjunctivitis

We identified a significant association for FEK (*P*=1.62×10-7) in the LMM GWAS, with a marker on chromosome E3, located 10.5 kb from the gene *TNFRSF17* (*tumor necrosis factor receptor superfamily, member 17*) (Fig. 3C). The second most significant association with this disease did not reach significance (*P*=3.1×10^−6^) but is located within the gene *TNFRSF21* (*tumor necrosis factor superfamily, member 21*). Both *TNFRSF17* and *TNFRSF21* belong to the tumor necrosis factor receptor superfamily, and *TNFRSF21* is expressed in the eye [24]. The E3 locus affects the risk for FEK in DSH cats (S3 Table).

#### IBD

A significant association (*P*=9.54×10^−8^) for IBD was identified using the FarmCPU GWAS. The marker is on chromosome B4 near the genes *ECHDC3 (enoyl-CoA hydratase domain containing 3)* and *USP6NL (ubiquitin-specific protease 6 N-terminal like)* (S2 Fig.). *ECHDC3* has a role in fatty acid biosynthesis and has been found to have an increased expression in the brains of Alzheimer’s patients [25] while *USP6NL* is a GTPase-activating protein for Rabs and is up-regulated in several cancers, including breast and colorectal cancers [26,27]. The B4 significant locus affects risk for IBD in DSH and DLH cats (S3 Table).

#### HCM

The LMM GWAS for HCM reached suggestive significance (*P*=2.76×10^−7^) with a marker on chromosome E3, located within the gene *SDK1* (*sidekick cell adhesion molecule 1*) (Fig. 4A), which is expressed especially in the kidney and retina [28,29] but has also been associated with hypertension [30]. This suggestive E3 locus affects risk for HCM in DSH cats (S3 Table).

**Figure 4.**
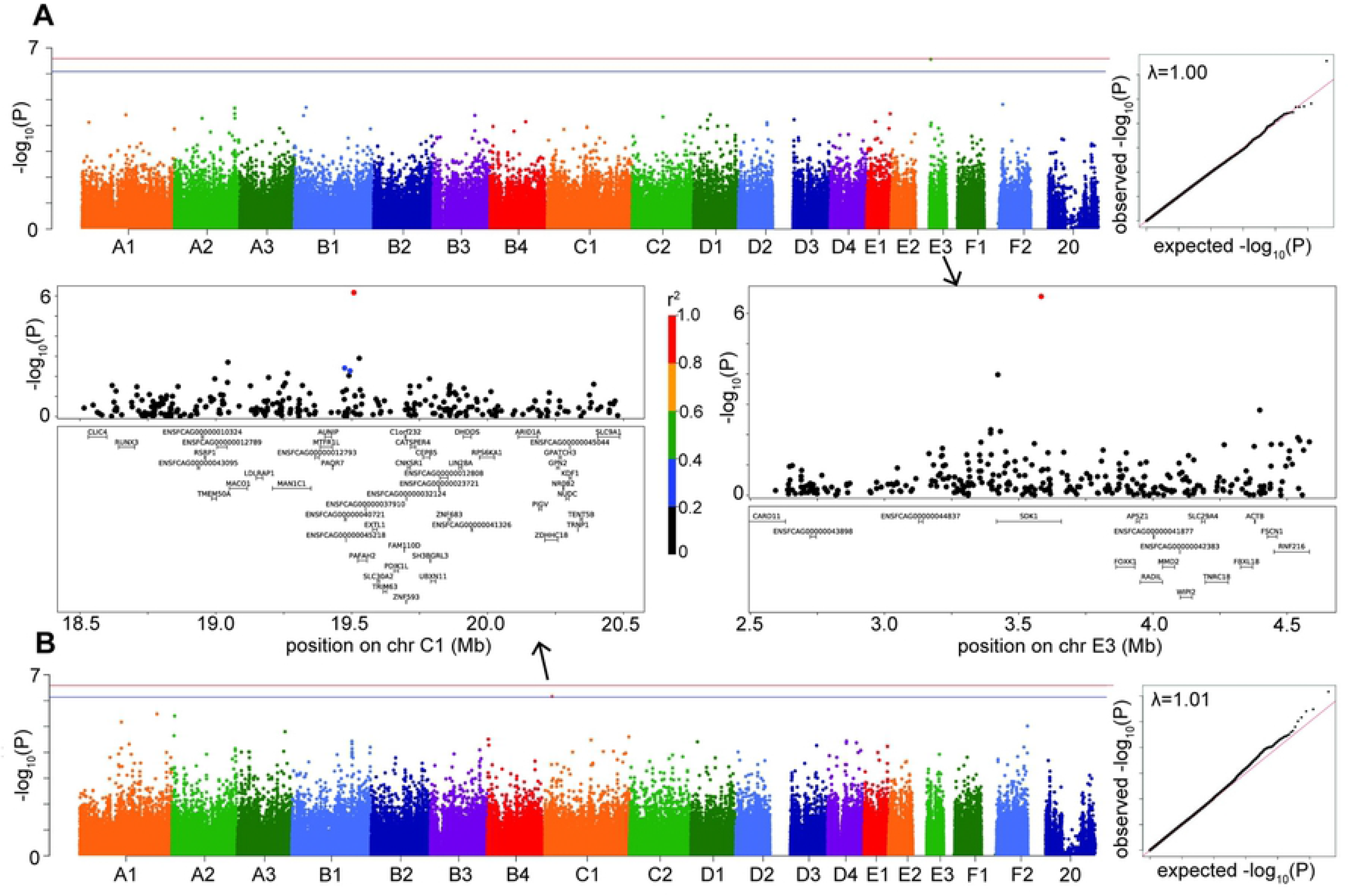
Manhattan, quantile-quantile (QQ), and LD plots for case-control disease suggestive associations, using the LMM GWAS results. X axis represents the chromosomal SNP position and Y axis represents the −log_10_(*P*-value). The QQ plots show observed versus expected *P*-values for each SNP. (A) HCM, showing the suggestive association on chr E3. (B) Hypercalcemia, showing the suggestive association on chr C1. On Manhattan plots, the red line is Bonferroni-corrected significance threshold, and the blue line is Bonferroni-corrected significance threshold calculated using unlinked SNPs. Inflation factors (***λ***) are shown on QQ plots. On LD plots, the colors indicate the amount of LD (r^2^) with the most significant SNP, ranging from black (r^2^<0.2) to red (r^2^>0.8).

#### Hypercalcemia

The hypercalcemia GWAS produced a suggestive association (*P*=6.81×10^−7^ in LMM, *P*=3.05×10^−7^ in FarmCPU) on chromosome C1, located in the gene *PAFAH2* (*Platelet Activating Factor Acetylhydrolase 2*) and within LD of the gene *STMN1* (*Stathmin 1*) (Fig. 4B). The enzyme encoded by *PAFAH2* acts to protect the cell from oxidative cytotoxicity [31], while the protein encoded by *STMN1* is involved in regulating the microtubule cytoskeleton, including mitotic spindle formation [32]. The C1 locus affects risk for hypercalcemia in DSH cats (S3 Table).

Genome-wide association studies of the other complex diseases, CKD, SCAL, chronic enteropathy, and merged GI phenotypes did not produce a significant or suggestive association using either the LMM or FarmCPU GWAS (S3 Fig., S2 Table, S4 Table).

## Discussion

In this study, we identified significant associations for common, clinically relevant, complex diseases in a population of 1,122 random and purebred cats, using a dense genotyping array. While a similar study was previously performed in dogs [33], this is the largest GWAS disease study in cats reported to date, conducted in a heterogeneous natural population including 80% random bred cats. Further advantages of the current study design were the careful phenotyping of aged control cats, accurate phenotyping of diseased participants by specialists performed in an academic clinical setting, and a mapping array approximately 5-fold denser than the current 63k array. Additionally, the quality of the biospecimens used and its associated data demonstrate the importance of using an accredited resource such as the Cornell Veterinary Biobank.

As a positive control, we identified significant associations for the *Orange* coat color locus and factor XII deficiency at the *F12* gene locus. Although the *F12* locus was the most significant association using both LMM and FarmCPU models, three and five other significant SNPs were identified in the factor XII deficiency GWAS, respectively. A BLAT [34] search showed that the flanking region of the three SNP maps to many places in the feline genome, including chromosome A1: 175 Mb, the location of the gene *F12.* Thus, it appears that there may be some non-specific binding with the A1 probe. However, factor XII deficiency is affected by several different loci across the genome, as shown in humans [35].

The majority of the cats included in our analyses are random bred cats, which generally have shorter LD than purebreds [3], because they have not been subject to selective breeding for specific traits. Further, we are mapping complex diseases, which usually consist of many variants each contributing a small effect, and have not been subjected to artificial selection, resulting in shorter LD surrounding the causal variant. As a result of investigating complex diseases in a predominantly random bred cat population, we do not expect to see the stacking of SNPs that are seen in GWAS studies of morphological traits, especially in purebred cats.

Using a case/control approach, we performed GWAS with both a LMM and FarmCPU, and found very similar results. Both methods identified significant associations for hyperthyroidism and DM, and the FEK association was significant in the LMM GWAS while the IBD association was significant in the FarmCPU GWAS. Furthermore, the same SNPs were identified as the most significant associations by both models.

For hyperthyroidism, the candidate gene *ARG1* encodes Arginase 1, a cytosolic enzyme that participates in the urea cycle and is expressed in the liver [36]. Another nearby candidate gene, *MED23,* encodes a component of the thyroid hormone receptor (TR) associated protein complex. As such, it interacts with, and facilitates, TR function. Variants in the TR have been associated with thyroid hormone resistance, for which the clinical presentation is very similar to thyrotoxicosis [37]. This is the first GWAS for feline hyperthyroidism reported and our finding represents a novel locus. Somatic variants in the *thyroid-stimulating hormone receptor (TSHR)* gene have been previously reported, but those variants were identified in DNA extracted from the affected thyroid glands of hyperthyroid cats [38].

The significant DM locus includes many olfactory receptor genes. Genetic and epigenetic variation, and the resulting functional changes, in olfactory receptors have been associated with taste, food intake, and satiety [39,40]. These differences may contribute to obesity risk and risk of DM. Mouse olfactory receptor gene OLFR15 has been shown to be expressed in pancreatic beta-cells and to regulate the secretion of insulin [41]. The other interesting gene near our significant D4 association, although not quite within the LD region of interest, *PTGS1,* encodes an enzyme that converts arachinodate into prostaglandin, which is involved in glucose homeostasis [42]. This gene has been associated with human DM [43,44]. Our findings constitute a novel locus associated with DM. Previous studies have identified several loci associated with DM in Australian Burmese cats [15,45] and a polymorphism in *melanocortin receptor 4 (MCR4)* associated with DM in obese domestic shorthair cats [46].

For eosinophilic keratoconjunctivitis, we identified a significant association in the LMM GWAS near the gene *TNFRSF17*. This is especially promising, and warrants further investigation because of its role in the innate and adaptive immune response. In patients with allergic asthma, eosinophils infiltrate the bronchial wall and lumen, and the bronchial epithelium is often damaged [47]. These pathological findings are associated with aberrant T helper 2 (Th2) cell-mediated immune responses. Interleukin-5, which is produced by Th2 cells, and the chemokine eotaxin are key players for the proliferation, differentiation, activation and mobilization of eosinophils [48,49]. In knockout mice studies, NF-kappa-B, a transcription factor that is activated by the *TNFRSF17* and *TNFRSF21* genes, was found to play an important role in Th2 cell differentiation and is therefore required for induction of allergic airway inflammation [48,50]. Similar to knockout mice with allergic asthma, it is possible that animals affected with FEK have an abnormal NF-kappa-B activation due to defective expression of *TNFRSF17* and *TNFRSF21* genes, as suggested by the current GWAS study.

The IBD significant association, as identified by the FarmCPU method only, was located near the genes *ECHDC3* and *USP6NL.* Neither of these genes are good candidates for a gastroenteropathy phenotype.

The first of the two suggestive associations, HCM, was located in the gene *SDK1*. A polymorphism in *SDK1* was found to be associated with hypertension in a study of over 5,000 Japanese individuals [30] but the function of this gene related to hypertension has not been described. Finally, the LD region surrounding the suggestive association for hypercalcemia contained the genes *PAFAH2* and *STMN1*, neither of which have been associated with hypercalcemia previously.

Despite the use of a dense genotyping array, across-breed GWAS for CKD, SCAL, chronic enteropathy, and merged gastrointestinal phenotypes did not reach statistical genome-wide significance using either single-locus or multi-locus models. We believe that larger cohorts may be needed due to the genetic architecture of these diseases, especially chronic enteropathy for which we had fewer than 50 cases in the respective GWAS.

By not restricting our analyses to a single breed, we were able to include a relatively large sample size for some of our phenotypes, thereby increasing statistical power to identify significant loci. In a study of this kind, especially if the majority of cats are randomly bred, LD is shorter, resulting in smaller regions of interest and narrowing the list of potential candidate genes.

Nevertheless, for some other phenotypes, we had an unbalanced proportion of cases and controls. This is due to the fact that accumulation of samples takes a long time, in part because donating samples is an opt-in process in our hospital, and because of the difficulty of recruiting universal controls. It is an unanswered question how many samples are required for a robust across-breed, complex disease GWAS study in cats, but canine simulation studies indicate that 500-1000 cases and controls, plus a further increase in array marker density, would substantially increase loci discovery in dogs [33].

Follow-up analyses using an independent cohort of phenotyped cats are needed to validate the associations we identified in this group of genotyped cats. Further studies involving investigation of the regions surrounding the significant associations are needed to determine causal variants for these complex diseases. Use of the >300 whole genome sequences provided by the 99Lives Feline Genome Consortium will allow variant discovery within candidate genes in the intervals of interest. Finally, functional studies will be required to confirm causal variants.

In this research, we used an across-breed GWAS design with a ~5-fold denser genotyping array than currently available, to identify significant associations with important common feline complex diseases. We demonstrated that a well-curated, hospital-sourced population can be used effectively for mapping studies. We also demonstrated the benefit of such a dense mapping array, propelling the field of complex feline disease genetics forward. Further, these results can be used to develop new diagnostic tests to assist veterinarians in identifying diseases earlier and allowing the implementation of early preventative measures. Breeders could improve their practices by identifying cats with optimum genetic value and owners could make informed decisions regarding the health of cats. This is particularly important in this era of personalized medicine. The shared environment of cats and their owners further enhances the value of domestic cats as models of lifestyle disease common to both species.

## Materials and methods

### Banking biospecimens and associated data

The 1,122 feline biospecimens used for this project were selected from the Cornell Veterinary Biobank (CVB), a core resource at the Cornell University College of Veterinary Medicine, which has been collecting and processing whole blood samples from feline patients admitted to the Cornell University Hospital for Animals (CUHA) since 2006. Biospecimens from participants consented at our satellite clinic, the Cornell University Veterinary Specialist in Stamford, Connecticut, were also included.

Out of the 1,122 cats, 57 were recruited through the Senior Feline Health Screening program from 2014 to 2018. The program was created to build a biobank of DNA and associated clinical data from healthy senior cats to serve as universal controls for mapping studies. In order to participate, feline candidates had to be at least 9.5 years of age and in good health. Privately owned cats that participated in the screening had a general physical examination and were examined accordingly by board certified specialists: cardiac auscultation and echocardiogram, dental examination, body condition scoring, body mapping (used by oncologists to record any masses found), ocular examination, and an orthopedic examination. A complete blood count, serum chemistry panel, coagulation panel, feline immunodeficiency virus (FIV) and feline leukemia virus (FeLV) test, baseline serum thyroxine (T4) level, and urinalysis were performed.

### Sample Processing, Storage and Distribution

Samples were collected according to the Cornell University Institutional Animal Care and Use Committee (IACUC) protocol #2005-0151. Following owner informed consent, whole blood samples were collected in EDTA tubes and refrigerated at 4°C until DNA extraction. Formalin fixed, paraffin embedded (FFPE) scrolls of splenic tissue were acquired from a collaborating pathologist and used for DNA extraction when necessary. Genomic DNA was extracted from blood samples using a standard salt precipitation. Genomic DNA was extracted from FFPE samples using the E.Z.N.A. Tissue DNA kit (Omega Bio-Tek) following the manufacturer’s instructions. DNA concentration and purity were determined by spectrophotometry on a NanoDrop ND1000 (Thermo Scientific). DNA samples were stored at ≤−20°C until distribution for genotyping.

### Inclusion criteria

Participants with a disease of interest could simultaneously be used as controls for other traits/diseases, as long as these traits were ruled out. Phenotypes included cases and controls from any breed, unless specified. Controls were at least 9.5 years of age, while cases could be of any age. Numbers of purebred and random bred cats included as cases and controls for each GWAS are shown in Table 2 and numbers of individuals from each breed are shown in S5 Table. The distribution of cases and controls by age is shown for each phenotype in S4 Fig.

**Table 2:** Distribution of cases and controls that are purebred and random bred cats for each disease.

| **Disease** | **PUREBRED** | **RANDOM BRED<sup>^</sup>** | **Total number** |
| --- | --- | --- | --- |
|  | **Number cases, number controls** | **Number cases, number controls** | **cases, controls** |
| **HCM** | 14, 20 | 69, 33 | 83, 53 |
| **Hyperthyroidism** | 24, 38 | 286, 96 | 310, 134 |
| **DM** | 13, 62 | 54, 304 | 67, 366 |
| **CKD** | 29, 10 | 125, 52 | 154, 62 |
| **Chronic enteropathy** | 9, 10 | 32, 18 | 41, 28 |
| **IBD** | 9, 15 | 38, 18 | 47, 33 |
| **SCAL** | 12, 16 | 66, 18 | 78, 34 |
| **GI combined** | 33, 16 | 133, 18 | 166, 34 |
| **FEK** | 2, 13 | 13, 27 | 15, 40 |
| **Hypercalcemia** | 5, 78 | 20, 365 | 25, 443 |

<sup>^</sup> includes domestic shorthair (DSH), domestic longhair (DLH), domestic medium hair (DMH), as well as cats identified as breed mixes (e.g., Siamese mix)

#### Hypertrophic cardiomyopathy

HCM is the most common cardiac disease in cats, affecting around 15% of the feline population [51,52]. Similarly to humans, familial HCM has been described in purebred cats, and in Maine coon and Ragdolls is caused by mutations in myosin binding protein C gene (MYBPC3) [53]. Some Maine coon and Ragdolls cats develop HCM in the absence of this mutation, indicating that other mutations are yet to be identified [53]. Diagnosis was based on echocardiography. Phenotypic criteria for controls included normal left ventricular wall thickness measurements: left ventricular free wall (LVFW) and interventricular septum (IVS) in diastole ≤ 6 mm by M-mode (motion mode). Phenotypic criteria for cases included LVFW and IVS wall thickness >6 mm. Additionally, affected cats must have had normal baseline T4 and be normotensive and normally hydrated in order to rule out other causes of cardiac hypertrophy.

#### Hyperthyroidism

Hyperthyroidism is one of the most common endocrine disorders affecting senior cats. The disease most often results from benign adenomatous thyroid nodules similar to human toxic nodular goiter [54]. Hyperthyroidism is believed to be a multifactorial disease, with nutritional, environmental, and genetic factors postulated as interacting causes [54]. The diagnosis of cases and controls was based on the following criteria: control cats had low-normal thyroxine (T4; <3 ◻g/dL; normal range 2-5 ◻g/dL). Cases had T4 >5 ◻g/dL or normal T4 with increased free T4. Radioiodinated thyroid scan results confirming the diagnosis were recorded, if available.

#### Diabetes mellitus

DM is also one of the most common endocrine diseases of cats with the majority of the cats resembling Type 2 (adult onset) DM in humans. The disease is caused by a combination of decreased β-cell function, insulin resistance, and environmental and genetic factors [55]. Diagnosis of DM was based on the following criteria: control cats had blood glucose values <200 mg/dL (reference values: 71-182 mg/dL) and no glucosuria. Cases had elevated blood glucose (>250 mg/dL) and glucosuria in at least two consecutive visits. Also, fructosamine, if evaluated, had to be above normal range (174-294 μmol/L). Of the 67 cases in the GWAS, 39 had fructosamine tests and all had elevated levels. Fifty-three diabetic cases and 339 controls had body weight recorded. Although the body weights of cases were spread throughout the range of 1.8kg to 10kg, a greater proportion of cases (12 of 53, or 22.6%) had weights >7kg, compared to controls (17 of 339, or 5.0%) (S5 Fig.).

#### Chronic kidney disease

CKD is highly prevalent in both humans and cats with approximately 10% of cats >10 years of age reported to be affected. Cats with CKD experience a progressive loss of functional renal mass. CKD is considered a heterogeneous syndrome, rather than a single entity [56]. CKD was diagnosed by evaluating the level of blood urea nitrogen (BUN) and creatinine, in conjunction with the urine specific gravity (USG). Symmetric dimethylarginine (SDMA), a natural occurring indicator for kidney function, was measured in the blood of some cases to determine if early renal disease was occurring. The diagnosis was established according to the following criteria: controls cats had creatinine <1.6 mg/dL (normal range 0.6-2 mg/dL), BUN within normal range (16-36 mg/dL) and USG >1.035 (preferably performed on the same day as creatinine was measured). Cases had to be azotemic (elevated BUN and creatinine values) with concurrent isosthenuria (failure of the kidney to dilute or concentrate urine) and increased SDMA, diagnosed by a board-certified veterinary internist.

#### Chronic enteropathy/inflammatory bowel disease/small cell alimentary lymphoma

Chronic enteropathies, which include Inflammatory Bowel Disease (IBD) and Small Cell Alimentary Lymphoma (SCAL), are common forms of primary gastrointestinal disease in cats. Although the cause of feline IBD is unknown, it has been hypothesized that, similar to canines and humans, feline IBD is caused by several factors such as intestinal microbial imbalances, diet, and defects in the mucosal immune system [57]. SCAL is the most frequent digestive neoplasia in cats, accounting for 60-75% of gastrointestinal lymphoma cases [58].

For this study, cats were assigned as chronic enteropathy cases if gastrointestinal (GI) clinical signs such as chronic vomiting, diarrhea, or weight loss were present, non-GI causes of their clinical signs were excluded, thus highly suggestive of either IBD or SCAL, but no histologic diagnosis was performed. IBD and SCAL were considered separate diagnoses that required histological confirmation. Distinguishing between IBD and SCAL can be difficult, so in addition to histologic assessment, immunophenotyping and polymerase chain reaction (PCR) for antigen receptor rearrangements (PARR) were used in some cases to confirm the SCAL diagnosis. Phenotypic criteria for affected cats included persistent clinical GI signs and histopathology performed by a board-certified veterinary pathologist confirming either IBD or SCAL. Control cats were examined by a board-certified oncologist and had an absence of any GI signs. We performed a separate GWAS for each of IBD and SCAL, and then chronic enteropathy, which includes cats that were not formally diagnosed but could be either IBD or SCAL. Finally, we performed a GWAS including all GI cases in an attempt to increase statistical power, and since IBD, SCAL, and chronic enteropathy can be considered a different manifestation of the same disorder [59]. There is also evidence that IBD leads to SCAL [60].

#### Feline eosinophilic keratoconjunctivitis

FEK is a corneal/conjunctival disease characterized by vascularized white-to-pink plaques on the cornea and bulbar conjunctiva. In the majority of cats, previous corneal ulceration has been diagnosed and an association with feline herpesvirus type 1 (FHV-1) infection has been proposed [61]. The diagnosis of FEK was made according to the following criteria: affected cats had signs of the disease during ophthalmologic exam performed by a board-certified veterinary ophthalmologist, including proliferative vascularized lesions affecting peripheral corneal/bulbar conjunctiva and the presence of eosinophils in the ocular cytology. Control cats had a normal ophthalmologic exam.

#### Hypercalcemia

Hypercalcemia is a common condition of cats defined by an increase in both total and ionized serum calcium. It may be caused by many conditions such as neoplasia, renal failure, primary hyperparathyroidism, hypoadrenocorticism, ingestion of cholecalciferol-containing rodenticides, or granulomatous disease. In cats, hypercalcemia can also be idiopathic [62], which is the phenotype we are investigating here. The diagnosis of hypercalcemia was determined as follows: control cats had total serum calcium values within the normal range (9.1-10.9 mg/dL); affected cats had elevated total serum calcium and ionized calcium values (reference interval 1.11-1.38 mmol/L). Parathyroid hormone (PTH) and PTH related peptide (PTHrP) were recorded if available, and were used to differentiate between causes of hypercalcemia.

### Design of array

Genotyping was performed on an Illumina Infinium iSelect Custom BeadChip. These arrays contain 340,000 attempted beadtypes for genotyping single nucleotide polymorphisms selected across the entire cat genome, using feline genome assembly felCat5. Of the 340,000 markers included on the array, 297,034 (87%) provided a reliable call.

SNPs for the array were selected from whole genome sequencing of 6 genetically diverse female DSH cats. These 6 cats were sequenced on a HiSeq2500 (Illumina, San Diego, CA) to generate 100bp paired-end reads. Following GATK best practices pipeline [63], reads were mapped to the feline reference genome using BWA mem [64], then duplicate reads were tagged by PICARD MarkDuplicates, and indels were realigned and quality scores were recalibrated using GATK. Variants were called and filtered using GATK HaplotypeCaller and VCFtools [65]. The full list of variants was thinned randomly using PLINK and then protein-coding variants with moderate and high impact as defined by SnpEff [66] were added back in.

### Genotyping

In total, 1,200 feline DNA samples were genotyped on the Hill’s custom Illumina feline high density mapping array. Genotyping was performed in 11 batches, or plates, by Neogen GeneSeek Operations (Lincoln, NE). Raw data files were converted to PLINK format and quality control was performed in PLINK v1.9 (www.cog-genomics.org/plink/1.0/) [67,68].

### Quality control

Genotyping data from the 11 batches were merged together using PLINK’s --bmerge command and a sex check of all samples was performed using PLINK’s --check-sex command. Seventy samples were removed due to missingness >80%, including 53 samples from the same batch.

SNPs were converted to the genome assembly felCat9 [69] and SNPs with missingness >95% in the 1,130 cats were removed, leaving 252,987 SNPs. Eight cats were genotyped on two different plates each as internal controls. The SNPs that were discordant between these eight duplicates were identified and removed. Finally, duplicate samples were removed, leaving a dataset of 1,122 individuals and 251,978 SNPs for GWAS.

A Principal Component Analysis (PCA) was performed using the program EIGENSTRAT in the EIGENSOFT package [70]. For this, linked SNPs were pruned using PLINK’s --indep 50 5 2 command, leaving 91,556 SNPs. PCA was performed using all cats to look for batch effects, and all purebred cats to ensure individuals of the same breed clustered together. PCA was also performed using only the cats included in each phenotype to identify and remove outliers before GWAS analysis. An outlier is an individual that is located separately from the main cluster of cats on either the PC1 or PC2 axis. Further, in order to reduce the effects of genetically distinct individuals in our GWAS, we also removed any purebred cat that was located separately from the main cluster of random bred individuals on either PC1 or PC2.

For the DM and HCM phenotypes, a further two and 16 cats, respectively, were genotyped on the same 340k custom Illumina array by external coauthors (MEW and JAB, respectively). For these cats, the genotype files were merged with the sample set before the QC was performed, as described above. The genotype and phenotype data for all three of these datasets are available as PLINK files, and include the SNP information (chromosome, bp location, alleles).

### Genome-wide association study

Both a single-locus linear mixed model (LMM) and a multi-locus model were used to perform a GWAS for each disease phenotype. The LMM was performed in the program GEMMA v 0.98.1 [71], which includes a relatedness matrix as a random effect. The multi-locus method performed was FarmCPU (Fixed and random model Circulating Probability Unification) [72] run using rMVP [73] in R. FarmCPU is designed to help control for false positives by including associated markers as covariates, while also reducing false negatives by removing the confounding between the population structure and kinship and the markers to be tested. We used the default parameters, with a maximum of 10 iterations. For each phenotype, we included the relatedness matrix calculated by GEMMA and a covariate file consisting of the first four PCs from a PCA run on the genotypes of the cats included in the phenotype only.

For both models, the Wald test was used to calculate *P*-values, and the Bonferroni correction (p_genome_=0.05) was used to calculate the genome-wide significance threshold. A suggestive threshold was calculated using the Bonferroni correction on unlinked SNPs (pruned using the -- indep 50 5 2 option in PLINK).

For each phenotype, PCA outliers and related cats (pihat>0.40) were excluded. Single nucleotide polymorphisms with a minor allele frequency (MAF) <5% and a genotyping call rate < 90% were removed from each analysis. SNPs are provided in genome assembly felCat9.

Manhattan and quantile-quantile (QQ) plots were created using the package qqman [74] in R v4.0.2 [75]. Lambda values, as a quantification for genomic inflation, were calculated in R. Linkage disequilibrium plots were created using matplolib [76] in jupyter notebook [77].

### GWAS positive controls

As a positive control for the 340k array, we performed GWAS on the presence of orange fur. The *Orange* locus has been refined to a 1.5Mb region on the X chromosome, although the causal variant is unknown [16–18]. We used 211 random bred cats in the orange GWAS: 90 cats that had a coat color description of orange (including solid orange, orange and white, and orange tabby), and 121 cats that had a coat color description of black, brown or brown tabby.

We also performed a positive control GWAS of factor XII deficiency, a common hereditary coagulation factor deficiency in cats that does not cause a bleeding diathesis. For this phenotype, affected cats were classified based on severe factor XII deficiency (factor XII coagulant activity < 10% of normal), whereas control cats had values above 60%. Nineteen affected cats and 34 controls were included in the GWAS.

## Acknowledgements

We especially thank the faculty and staff of the Cornell University Hospital for Animals for sample collection and phenotyping of control cats. We would like to acknowledge Gregory Acland and John Schimenti for instigation of the Cornell Veterinary Biobank, and support from Baker Institute for Animal Health, Cornell University Center for Vertebrate Genomics, and Department of Clinical Sciences. Finally, we thank numerous pet owners for donating their pets’ samples, and collaborators for sample collection and phenotyping.

## Supporting information

**S1 Fig. LD plots for the GWAS positive controls, using the LMM GWAS results.** (A) *Orange* locus GWAS, showing the significant association on chromosome X. (B) Factor XII deficiency GWAS, showing the A1 significant association in the vicinity of the gene *F12.* Colors indicate the amount of LD (r^2^) with the most significant SNP, ranging from black (r^2^<0.2) to red (r^2^>0.8).

**S2 Fig. Manhattan, QQ and LD plots for IBD using the FarmCPU GWAS results.** On the Manhattan plot, the red dashed line is the Bonferroni-corrected significance threshold. Inflation factor (***λ***) is shown on the QQ plot.

**S3 Fig. Manhattan and QQ plots for diseases that did not reach genome-wide significance in the LMM GWAS.** On Manhattan plots, the red line is Bonferroni-corrected significance threshold, and the blue line is Bonferroni-corrected significance threshold calculated using unlinked SNPs. Inflation factors (***λ***) are shown on QQ plots.

**S4 Fig. Age distribution of cases and controls.** (A) HCM, (B) hyperthyroidism, (C) DM, (D) CKD, (E) chronic enteropathy, (F) IBD, (G) SCAL, (H) all GI, (I) FEK, (J) hypercalcemia. Age (in months) is shown on the X axis and number of cats is shown on the Y axis. Cases are the pink bars and controls are the blue bars.

**S5 Fig. Weight distribution of cases and controls for DM.** Weight (in kg) is shown on the X axis and number of cats is shown on the Y axis. Cases are the pink bars and controls are the blue bars. Note that weight data was only available for 53 of the 67 cases, and 339 of the 366 controls used in the GWAS.

**S1 Table. Significant association LMM GWAS results on chromosome X for the presence of orange fur in random bred cats.**

**S2 Table. Three most significant associations for each disease from the FarmCPU GWAS results.**

**S3 Table. Frequencies, odds ratios, and *P*-values for SNPs associated with complex diseases in different breeds and random bred cats.**

**S4 Table. Non-significant associations identified for complex diseases using a LMM GWAS design.**

**S5 Table. Numbers of individual cats from each breed and random bred group for each phenotype.**

